# Thermal Fogging with Disinfectant Didecyl Dimethyl Ammonium Bromide Effectively Kills A Coronavirus, An Influenza Virus and Two Indicator Bacteria in Subzero Cold-Chain Environment

**DOI:** 10.1101/2021.03.25.436894

**Authors:** Qiaoyun Hu, Pei Ma, Junyi Hong, Yulong Wang, Dong Huang, Yadi Tan, Zhengjun Yu

## Abstract

**Background:** The origin of several local COVID-19 outbreaks in China in 2020 were confirmed to be frozen food or packages contaminated with SARS-CoV-2, revealing the lack of effective disinfection measures in the frozen food chain.

**Objective:** To evaluate the disinfection efficacy at −20°C of a recently marketed thermal fogging disinfection product, with disinfectant-antifreeze combination being didecyl dimethyl ammonium bromide (DDAB) - propylene glycol (PPG).

**Method:** Carriers with porcine epidemic diarrhea virus (PEDV) YN1, a coronavirus, and a swine influenza virus H1N1 (SIV-H1N1), and three indicator bacteria, *E. coli, S. aureus,* and *B. subtilis* endospores, were respectively placed in a −20°C freezer warehouse with or without DDAB-PPG fogging and activities of the microorganisms were tested.

**Results:** At −20°C, DDAB-PPG fogging, which fully settles in 3.5−4.5 hours, fully inactivated PEDV of 10^−3.5^ TCID_50_/0.1ml and SIV-H1N1 of 2^6^ hemagglutination titer within 15-30 min, and inactivated *S. aureus* and *E. coli* vegetative cells (10^6^/ml) within 15 or 60 min, respectively, but had little effect on *B. subtilis* spores. The bactericidal effect lasted at least 3 hours for bacteria on carrier plates and for 6 hours for airborne bacteria.

**Conclusions:** A practical subzero temperature disinfection technology was confirmed its efficacy in killing enveloped viruses and vegetative bacteria. It would help to meet the urgent public health need of environmental disinfection in frozen food logistics against pandemic and other potential pathogens and to enhance national and international biosecurity.

**Highlights:**

- First report of efficacy of disinfectant didecyl dimethyl ammonium bromide (DDAB) at subzero temperatures
- The customized thermal fogging machine makes fine droplets of DDAB-propylene glycol (PPG) fog which can suspend for ~4 hours at −20°C
- DDAB-PPG thermal fogging at −20°C effectively inactivated a coronavirus and an influenza virus
- DDAB-PPG thermal fogging at −20°C for effectively killed *S. aureus* and *E. coli*
- DDAB-PPG thermal fogging at −20°C inactivated airborne microorganism for up to 6 h

## 1. Introduction

The objective of this study is to evaluate the disinfection efficacy of a thermal fogging method in a temperature for frozen foods. This is an urgent market need because SARS-CoV-2 present in frozen food ^1^ or packages ^2^ has been identified as the origin of transmission in some recent local outbreaks of COVID-19 in China.

Considering that high SARS-CoV-2 infection rate was found among meat processing plant workers^3^ and fishery workers ^4^ early in the pandemic, and SARS-CoV-2 can survive for a long time under cold temperatures ^5,6^, these constitute a full cycle of human-to-frozen food-to-human transmission of SARS-CoV-2, which reveal both an important public health threat and a weakness in the disinfection procedures of the frozen food cold-chain.

As frozen temperature is inhibitory for growth of bacteria and viruses, and most food cold-chain related viral pathogens impact the gastrointestinal system and are often linked to incomplete cooking of food ^7^, and none about respiratory diseases. Routine disinfection practice has been insufficient if not absent in some critical links of the frozen food cold-chain operations. But the new finding that human respiratory viruses are transmitted via frozen food calls for a safer cold-chain with effective disinfection measures ^8^.

Facing the situation, China issued a number of technical guidances on food cold-chain disinfection ^9^, including *Technical guidance for prevention and control of COVID-19 in cold food production chain* and *Technical guidance for disinfection in cold food production chain to prevent and control COVID-19*” Guidance [2020]-245 (Oct 16, 2020) ^10^ and “*Work plan for preventive and comprehensive disinfection of imported cold-chain food*” Guidance [2020]-255 (Nov 8, 2020) ^11^. A key focus of these guidelines is preventive disinfection of food product packaging and interior surfaces of shipping containers, particularly in the frozen food import chain from port to market, instead of food content within packaging. These guidelines require disinfection be carried out at critical steps and efficacy of disinfection checked, and call for “sum up good experience and practice in SARS-CoV-2 detection and disinfection” instead of giving specifics on how to disinfect, particularly for the frozen chain. Thus it can be said that the disinfection practices in the frozen food chain are still being explored and optimized. Many disinfection products have been used since the COVID-19 pandemic, but data are needed to support their efficacy and continued use in the frozen food cold-chain.

Therefore, we carried out disinfection efficacy study of a recently marketed disinfection product suitable for freezing temperatures. It uses a customized thermal fogging machine to deliver a disinfectant solution together with a matching fogging fluid & antifreeze. The disinfectant is didecyl dimethyl ammonium bromide (DDAB), and the fogging fluid & antifreeze is propylene glycol (PPG). DDAB is among the most widely used disinfectants worldwide and is on the recommended cold-chain disinfectant list by health authorities. Two field strain viruses from the coronavirus family and influenza virus family, and three indicator bacteria, *E. coli, S. aureus,* and *B. subtilis* endospore were tested in a −20°C food industry freezer warehouse. The data would support informed choices for cold-chain disinfection practice.

## 2. Material and Methods

### 2.1 Indicator bacteria

The *Escherichia coli* strain ATCC8099, *Staphylococcus aureus* strain ATCC6538 *Bacillus subtilis* spores (Guangdong Microbial Culture Collection or GDMCC1.372) were used.

### 2.2 Viruses

The porcine epidemic diarrhea virus (PEDV) field strain was a gift from Dr. Qigai He of Huazhong Agriculture University, and its passage and Median tissue culture infective dose (TCID_50)_ assay were carried out in Vero cells^12^. The swine influenza virus (SIV) H1N1 A/S/TJ/04 strain was a gift from Dr. Meilin Jin of Huazhong Agriculture University and its passage and infectivity titer determination were carried out on embryonated eggs^13^.

### 2.3 Disinfectants and thermal fogging at −20°C

The Huanying 360 thermal fogging machine uses a didecyl dimethyl ammonium bromide (DDAB) water solution of 50g/100 ml and a 50% (v/v) propylene glycol (PPG) fogging and anti-freeze fluid with 0.1% lemon fragrance limonene. The combination is referred as HY-360 DDAB-PPG. The machine is programmable for duration of fogging. After setting up and powered on, it runs and stops automatically. The concentration of DDAB-PPG fog used in the study are listed in Table 1, which included two concentrations, ~0.4 g/m^3^ and ~1.0 g/m^3^, corresponding to ~8 or ~20 min of fogging. A photo of the fogging machine in operation is shown in Figure 1.

**Table 1.** Volume of the −20°C freezer warehouses and fogging doses

| Disinfection experiment | Figure | Volume (m<sup>3</sup>) | DDAB used (ml) | PPG used (ml) | DDAB (ml/m<sup>3</sup>) | DDAB (g/m<sup>3</sup>) |
| --- | --- | --- | --- | --- | --- | --- |
| PEDV in 1 h | Figure 3 | 168 | 330 | 400 | 1.96 | 0.98 |
| SIV in 1 h | Figure 3 | 97.44 | 80 | 120 | 0.82 | 0.41 |
| 3 bacteria in 1 h | Figure 4 | 97.44 | 80 | 110 | 0.82 | 0.41 |
| 3 bacteria in 1 h | Figure 4 | 273.83 | 550 | 600 | 2.01 | 1.00 |
| bacteria in 8 h | Figure 5 | 273.83 | 550 | 600 | 2.01 | 1.00 |

**Figure 1.**
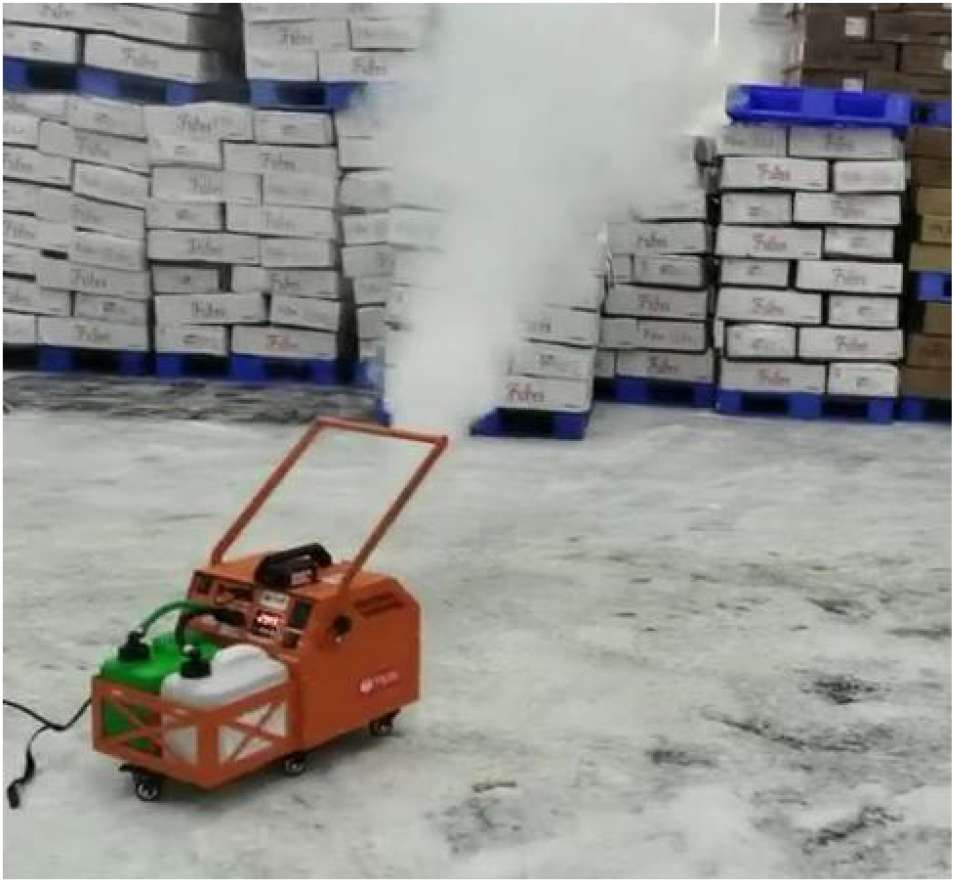
A photo of Huanying 360-DDAB thermal fogging machine (lower right) started fogging operation in a freezer warehouse (This is from a video of fogging which can be provided as supplementary data).

### 2.4 Evaluating the efficacy of disinfectant fogging on PEDV at −20°C

The PEDV virus stock was prepared to be 10^−3.5^ TCID_50_/0.1ml. Carrier polystyrene plates were prepared by dropping 60 μl virus stock in each clean plate and let air dry, which were placed either on the floor or on a shelf in the freezer warehouse and exposed to disinfection thermal fogging for 15-60 min as indicated. Some plates served as freeze-only controls. The plates were taken out, rinsed with 1.2 ml DMEM (containing 2% penicillin and streptomycin), and the eluents were collected and stored at −80C freezer for TCID_50_ assay later, which is done according to previous described method.

### 2.5 Evaluating the efficacy of disinfectant fogging on SIV H1N1 at −20°C

SIV H1N1 stock of HA titer 1 : 2^5^ was diluted 2 fold with sterile saline, polystyrene plates were prepared by dropping 50 μl virus stock in each plate and let air dry. The carrier plates were placed on different locations in the freezer warehouse as indicated and exposed to disinfectant thermal fogging for 15-60 min as indicated. Some plates were served as freeze-only controls. The plates were taken out, each was rinsed with 1.5 ml sterile saline, and the eluents were inoculated at 0.4 ml/each onto three 10-day chicken embryos and cultured at 37℃ for 72 h. Then the eggs were put into −20℃ for 1 h, and the allantoic fluids were collected, 2 fold serial diluted with saline and used for hemagglutination assay with chicken erythrocytes. The highest dilution with erythrocytes agglutination was recorded as the rough HA titer of that sample. To make titers from different assay runs comparable, the mean titer of the stock assayed in each run was divided by 2^5^ to obtain a normalization number and all the titers obtained were divided by this number.

### 2.6 Bacteria disinfection experiments at −20°C

*E. coli,* and *S. aureus* were cultured in 10-20 ml TSB (Soybean-casein-digest) medium overnight in 50 ml conical tubes, counted, and adjusted to to 10^6^ cfu/ml. For *B. subtilis* spores, dry powder were resuspended with 0.85% saline, counted, and then diluted to 10^6^ cfu/ml saline. Then 100 μl each was spread on a 60 mm diameter empty PS plate, let air fry in biosafety hood and then covered. Carrier plates were placed in a −20°C freezer warehouse either on the floor (2 sites) or on a shelf (1.2 m height, 2 sites) at 2-3 plates each site, and exposed to disinfection fog for 15-60 min as indicated.

For evaluating the disinfection effect on airborne bacteria, clean plates were placed into the freezer warehouse at specified time, uncovered for 15 minutes to collect bacteria in the air, then covered, taken to the lab, rinsed with 1.0 ml sterile saline. The eluents were spread onto fresh sterile TSA-agar plates at 100 μl per plate and cultured overnight at 37°C for colonies counting to obtain cloning forming units (cfu).

### 2.7 Data analysis

The disinfection efficacy of DDAB fog for PEDV and the 3 bacteria were assessed by calculating the log_10_ TCID_50_/0.1ml or cfu and comparing fog exposed samples to the starting value or freezer only controls. Reduction of >=3log_10_ is considered qualified. For SIV H1N1, the infectivity is measured by HA titer which is expressed as log_2_ values. The figures were plotted with Plot2 Pro software.

## 3. Results

### 3.1 The DDAB-PPG fog produced by HY360 thermal fogging machine can suspend for several hours in the air

A series of photos were taken for the process of disinfection fogging with distance markers present (shown in Figure 2). The visibility at time 0, i.e, right after the machine finished fogging, was 0 at all conditions tested. It took ~3.5 h or ~4.5 h for the fogged rooms to restore visibility to > 5 meters at room temperature (RT) or −20°C, respectively.

**Figure 2.**
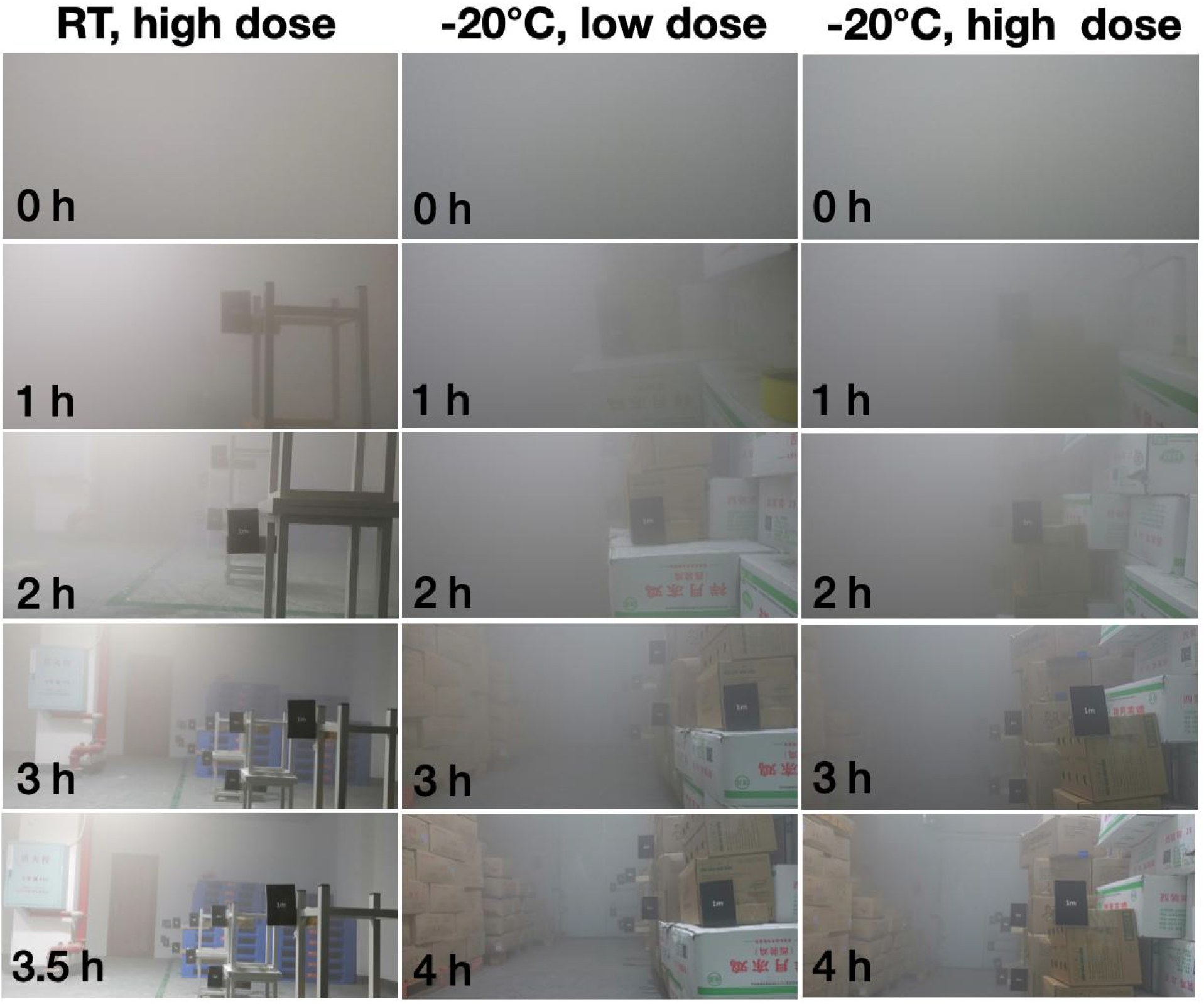
Photos of HY-360 DDAB-PPG thermal fogging in a RT room or a −20°C freezer warehouse. The high dose is ~1 g DDAB/m^3^, low dose is ~0.4 g DDAB/m^3^. Time 0 h is when the machine finished fogging.

The long hours of suspension gives the disinfectant fog ample time to act on all surfaces exposed and arrive hard-to-reach spots. In addition, based on the settling speed (5 meter high/~3.5 h at RT) the fog droplet size can be estimated by Stokes law, which should give the droplet size around 4~6 μm^14^.

### 3.2 DDAB thermal fogging has satisfactory virucidal efficacy at −20°C on PEDV and SIV-H1N1

Thermal fogging with ~1g/m^3^ DDAB at −20°C completely inactivated PEDV with log_10_ TCID_50_ titer of 3.5, and SIV H1N1 with HA titer of 2^5^. The experiment was repeated with 0.4 g/m^3^ DDAB for SIV H1N1, and full inactivation was achieved in all expect in 1 out of 6 sample at 15 min (Figure 3).

**Figure 3.**
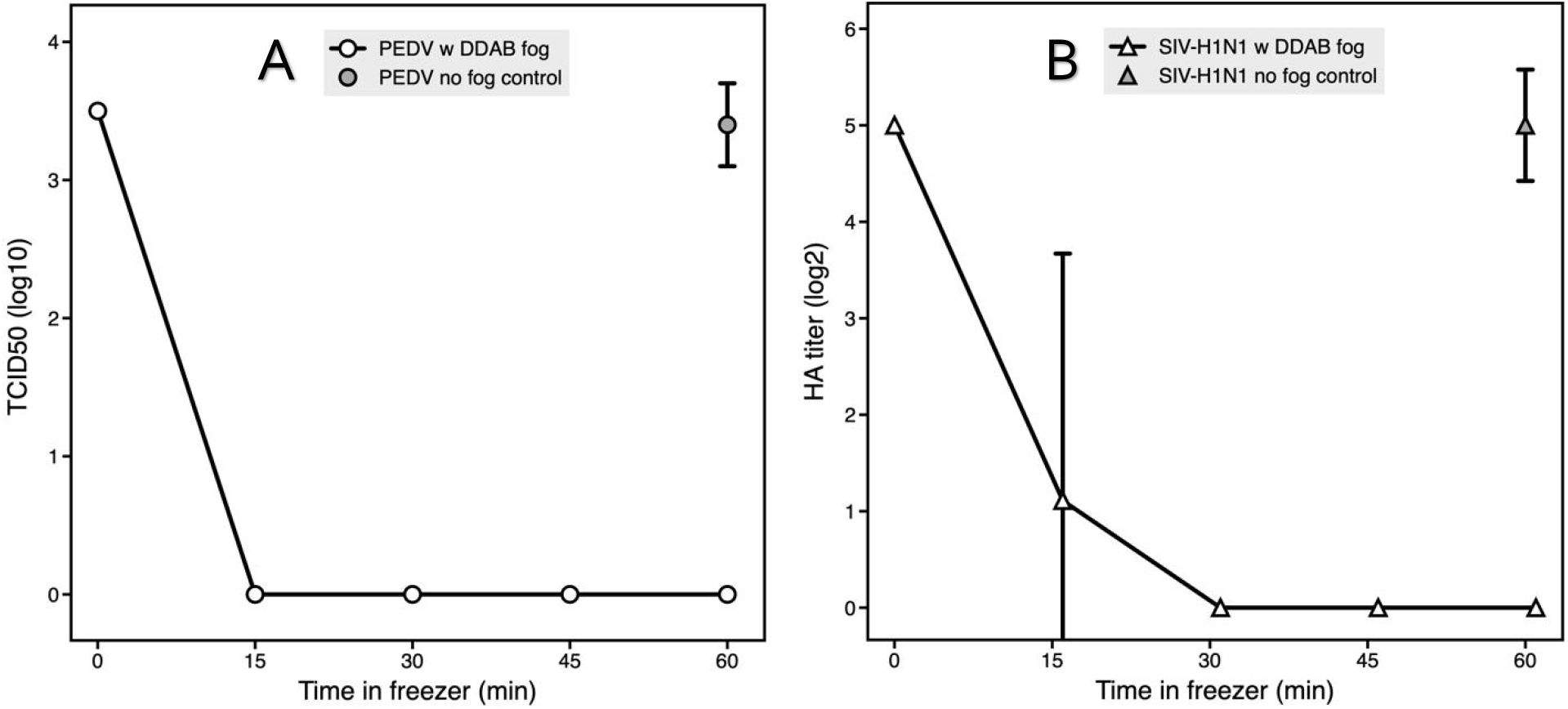
The virucidal effect of DDAB thermal fogging in a −20°C freezer warehouse. The fogging dose shown here was 1 g/m^3^ for PEDV (A) and 0.4 g/m^3^ for SIV H1N1(B). Data for 1g/m^3^ fogging for SIV H1N1 were all 0 after 0 min and not shown. For each data points n=8 for PEDV and n=6 for SIV H1N1.

### 3.3 DDAB thermal fogging at −20°C effectively disinfected vegetative bacteria cells but not spores

Thermal fogging with 1.0 g or 0.4g/m^3^ DDAB in a −20°C freezer warehouse produced ~5 log_10_ reduction of bacteria *S. aureus* within 15 min for both doses, and >4 log_10_ reduction of *E. coli* within 30 min or 60 min for the 1g and 0.4 g dose, respectively; as controls, the 60 min freezer exposure only samples had only <1.5 log_10_ reduction. However, with *B. subtilis* spores, DDAB thermal fogging only resulted in 1-2 log_10_ count reduction while freezer only control had similar count (Figure 4).

**Figure 4.**
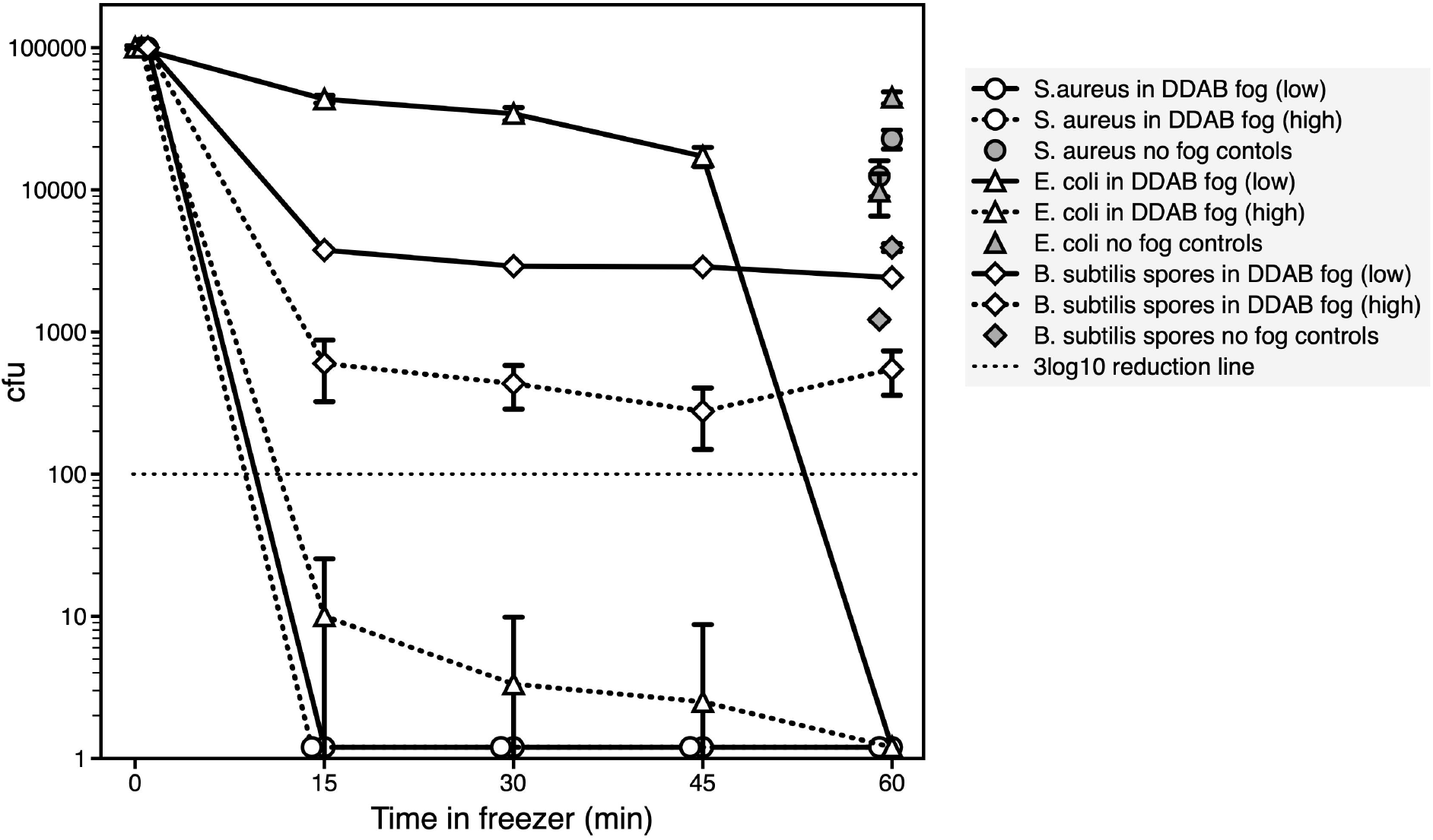
The bactericidal effect of DDAB fogging in a −20°C freezer warehouse. Each carrier plate was prepared with 10^5^ bacteria vegetative cells (*S. aureus and E. coli*) or spores (*B. subtilis*) at time 0. Then placed in the freezer with or without fogging. n=12 for each data point. For cfu of 0, a value of 1.2 was assigned to be plot on the log scale plot.

### 3.4 The disinfection effect of DDAB thermal fogging lasts for 3-6 hours

One time DDAB thermal fogging showed 3 or 5 hours of lasting effect (>3 log_10_ reduction) on *S. aureus* and *E. coli* bacteria, respectively (Fig.5A) and can have up to 6 hours of effect in eliminating airborne bacteria in the freezer warehouse (Fig. 5B).

**Figure 5.**
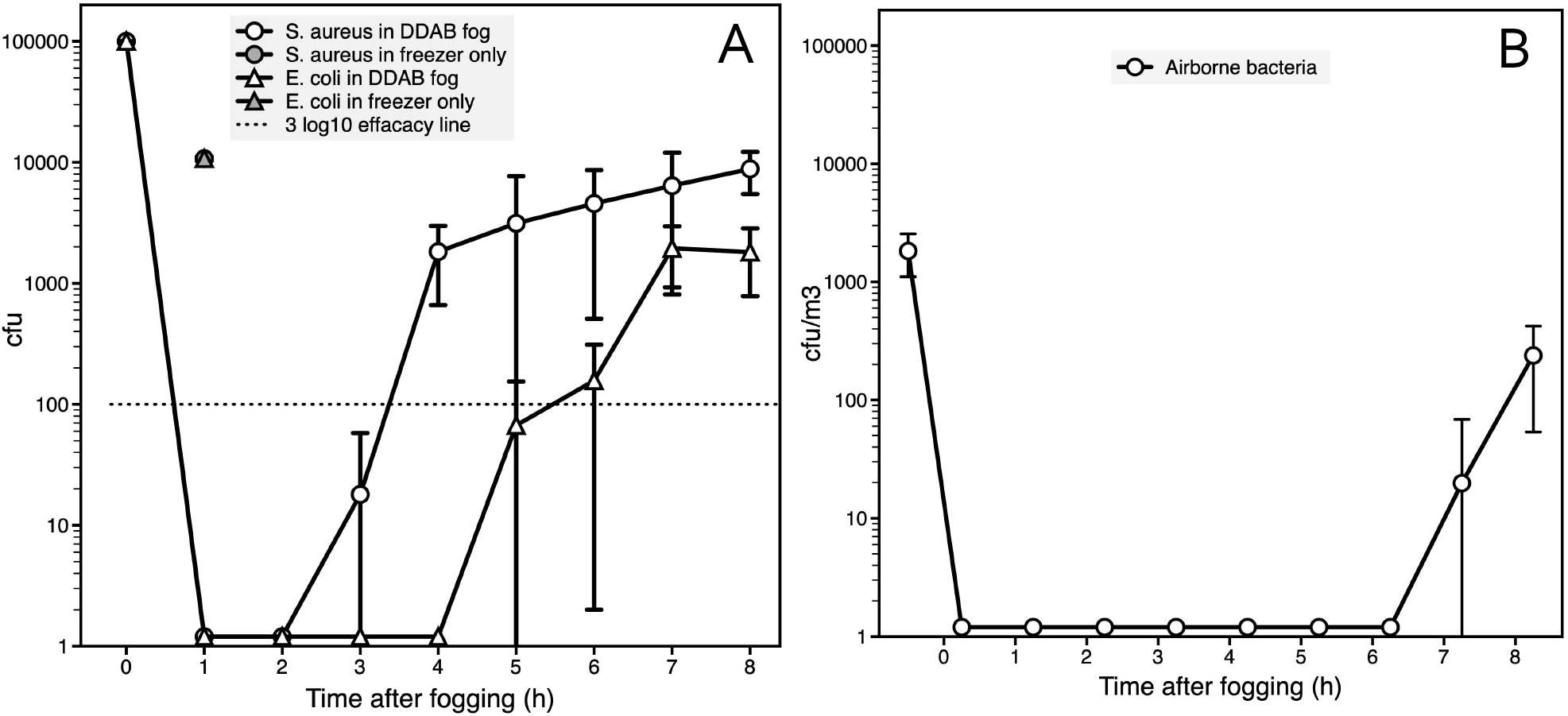
The lasting bactericidal effect of DDAB fogging at −20°C. The x value corresponds to time of sample taking out of freezer. In A, the carrier plates were exposed for 1 h (n=6); in B, the sterile plates were exposed for 15 min (n=6).

## 4 Discussions

The significance of the innovation reported is that it meets practical needs of disinfection in the food cold-chain industry under COVID-19 pandemic.

We here provide evidence that a practical innovation in disinfection based on optimizing existing technology could meet a global unmet need in subzero cold-chain disinfection, which can help to break the chain of SARS-CoV-2 transmission around the globe via cold chain so as to prevent re-emergence of the disease in areas already under control, and could also help to minimize the transmission of other human pathogens or bioterrorism agents though the cold-chain.

I DDAB is well known to kill viruses and bacteria by membrane disruption. However, its use in subzero temperatures had not been reported before. As it has higher efficiency at room temperature than at 4°C at inactivating viruses and vegetative bacteria^15^, it is necessary to test its disinfection efficacy at freezing temperatures, which means it first has to be delivered in a form and dose that do not freeze and suitable for frozen food cold-chain.

Disinfection with DDAB in subzero temperature was achieved by tailor-made optimization of thermal fogging technology. Although thermal fogging technology has been widely used in plant protection, vector control, disinfection and sanitation for decades, the technology usually requires optimization for specific purposes, both in terms of machine engineering and chemical selection which is related to the active agent to be delivered, such as pesticide or disinfectant, and the range of the fog to reach. For example, a study of atomizer optimization found that structural parameters of the atomizer of a fogging machine is key for the size of fog droplets, and with proper engineering, fog droplets could be controlled within a narrow range of 4.5~6.5 μm^16^. The fogging fluid, or solvent, is also key for fogging features including fog volume and droplet size. The thermal fogging machine Huanying360 is designed and optimized specifically for DDAB fogging, the combination haunting360-DDAB-PPG, the thermal fogging machine plus disinfectant DDAB and fogging fluid propylene glycol, studied here was the best outcome from many earlier explorations.

To our knowledge, this is a first report of efficacy of disinfectant DDAB at subzero temperatures, regardless of bacteria or viruses. The two viruses tested, a coronavirus and an influenza virus, are within the two most important virus families that have caused pandemics. The virucidal and bactericidal efficacy of DDAB-PPG thermal fogging shown in figures 3–5 strongly support the effectiveness of the routine dose (~0.4 g/m^3^) of DDAB-PPG thermal fogging for regular disinfection of surfaces in frozen food logistics. The dose can be doubled to achieve stronger and faster effect if heavy contamination is suspected.

The advantages of the technology presented in this study are summarized below.

First, thermal fogging technology disperses the disinfectants throughout the air and into hard-to-reach spots to achieve dead-zone free disinfection of exposed surfaces of all angles, and it is likely to be able to eliminate all airborne viruses as the fine fog droplets permeate the whole space for hours.

Second, the product uses familiar chemicals with excellent safety profile.

The disinfectant chosen, DDAB, is officially approved and recommended in recent disinfection guidelines for SARS-CoV-2 and had many years of use in numerous products. The fogging fluid, PPG, which also function as antifreeze, is often used to make artificial smoke and mists for fire safety training, theatrical performances and rock concerts, and has been recognized as safe^17^.

Third, minimal human labor is required for using the machine-reagent product set.

The reagents are ready to use and no dilution with water is required which reduces workload and chances for errors. Once set and powered on, the machine will run and stop automatically. As seen in Figure 1 and 2, the fogging projects at least 5 meters high and fogging from a single spot can fill a room of ~300 m^3^ by diffusion. For moderate spaces such as a shipping container or a truck, usually one machine operating from one spot is sufficient.

Fourth, it is economic for regular disinfection in industrial use, as has been proven in room temperature settings as compared to other options.

Regarding limitations of the technology, repeated fogging without cleaning could result in accumulation of antifreeze PPG which at high concentrations is slippery, therefore anti-slippery measures should be kept in mind in operation. In addition, the fogging machine is electrically powered, and therefore needs power supply to run, which is not as mobile as diesel fueled thermal fogging machines. Different thermal fogging machines can be chosen based on practical purposes and requirements.

## 5. Conclusions

Effective and practical subzero cold-chain disinfection of human viruses and other potential pathogens is an urgent and important public health and biosecurity need of the globe at the present and into the future. The HY360-DDAB-PPG fogging product reported in this study can meet this need with efficacy, safety, economy and practicality.

## CRediT authorship contribution statement

**Qiaoyun Hu:** Conceptualization, Methodology, Investigation, Data Curation, Writing - Review & Editing.

**Pei Ma:** Investigation, Supervision, Resources, Data Curation, Writing - Original Draft.

**Yulong Wang:** Investigation, Methodology, Validation. **Dong Huang:**Investigation, Data Curation, Validation. **Junyi Hong:**Investigation. **Yadi Tan:**Data Curation, Writing - Original Draft, Writing - Review & Editing. **Zhengjun Yu:**Conceptualization, Funding acquisition, Resources, Supervision, Writing - Review & Editing.

## Appendix A. Supplementary data

Supplementary data is a 15 second video of thermal fogging disinfection in action in a freezer warehouse.

